# Pathogen-induced flavonoid–anthocyanin biosynthesis in *Chrysanthemum seticuspe* confers resistance to *Botrytis cinerea*

**DOI:** 10.64898/2025.12.20.695701

**Authors:** H.M. Suraj, Lisa-Marie Roijen, Frank P.J. Pieterse, Malleshaiah SharathKumar, Kumar Saurabh Singh, Justin J.J. van der Hooft, André Fleißner, Jan A.L. van Kan

## Abstract

*Botrytis cinerea* is a major threat to ornamental crops, yet floral defence responses remain poorly understood. In *Chrysanthemum seticuspe,* we found that flower petals, unlike leaves, mount a localized resistance response resulting in red spots appearing at fungal penetration sites. We observed an intensification of the coloured response as infections progressed. To investigate the basis of this phenotype, we performed a time-course paired transcriptomic and metabolomic analysis on mock-inoculated vs *B. cinerea-*inoculated petals and leaves. Infection triggered strong transcriptional reprogramming in petals, with clear induction of phenylpropanoid and flavonoid/anthocyanin pathway genes and candidate regulators, consistent with the visible pigmentation. Metabolite profiles reflected this response, showing time-dependent accumulation of infection induced flavonoids such as quercetin, tilianin, and their derivatives, as well as cyanidin-based anthocyanins in infected petals. Integrating both omics datasets with MEANtools highlighted an anthocyanin-associated transcript–metabolite module, including a module putatively involved in the synthesis of polyyne-type phytoalexins. Antifungal assays demonstrated that selected flavonoids and cyanidin derivatives inhibit *B. cinerea* in a dose-dependent manner, supporting a direct antifungal role of these compounds. Altogether, our results show that *C. seticuspe* petals deploy a spatially confined, multi-layered chemical defence in which pathogen-induced flavonoids and anthocyanins operate as active components of resistance against a necrotrophic pathogen. We anticipate that future paired omics analyses in combination with spatial omics and bioactivity assays will yield insights into the role of specialised defence molecules in response to biotic and abiotic stresses.

**Graphical abstract:** 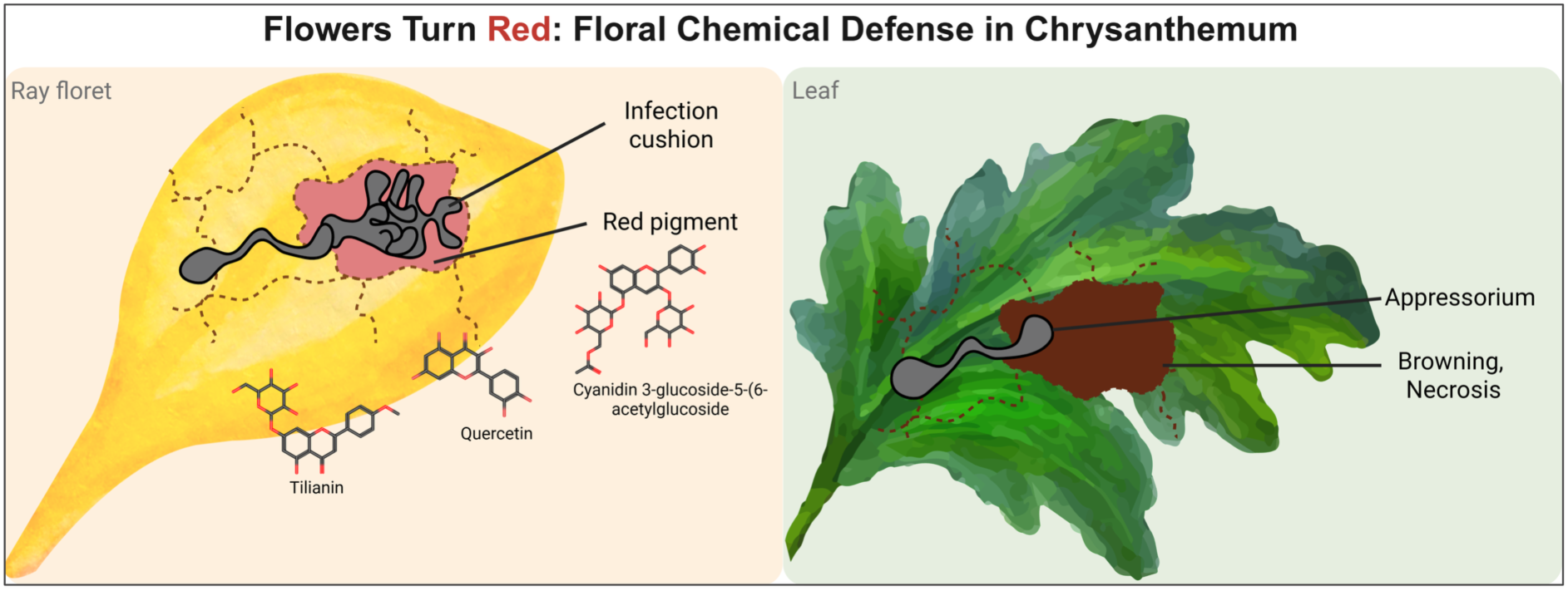

## Introduction

Plants rely on chemical defence mechanisms to counter pathogen attack. A hallmark of these defences is the rapid, localized accumulation of specialized (“secondary”) metabolites at the infection sites. In most cases, infection results in browning or clearing of plant tissue but, in some species, the responses to infection are colourful. Among such metabolites, flavonoids including anthocyanins and flavonols play diverse roles in plant survival, contributing to defence against pests and pathogens, UV protection, and pigmentation that deters herbivores and attracts pollinators (Agati et al., 2025; Agati & Tattini, 2010; Buer et al., 2010; Glover & Martin, 2012; Koes et al., 2005; L. Wang et al., 2022; Zhang, Butelli, De Stefano, et al., 2013). For example, tea leaves inoculated with *Colletotrichum camelliae* develop pink rings around lesions due to accumulation of pathogen-induced anthocyanins which functions as phytoalexins (Li et al., 2023).

Plants synthesize a broad spectrum of specialised metabolites, including terpenoids, alkaloids, and phenolics, as part of their defence repertoire. Within this chemical arsenal, flavonoids stand out due to their multifunctionality and protect plants from both biotic and abiotic stress. Increasing evidence shows that flavonoids, including anthocyanins, contribute to plant defence across diverse plant–pathogen systems. They can act directly against pathogens by disrupting fungal membranes or inhibiting pathogen enzyme activity, while also modulating host defence signalling. In addition, anthocyanins can scavenge reactive oxygen species (ROS), limiting oxidative damage and supporting cellular homeostasis in the host plant during infection (Agati et al., 2025; Agati & Tattini, 2010; Daryanavard et al., 2023; Wang et al., 2022). Flavonoid biosynthesis is tightly regulated by the conserved MYB–bHLH–WD40 (MBW) transcriptional complex (S. Li, 2014), which activates biosynthetic genes to enable rapid pigment accumulation at infection sites. However, despite these insights, the precise contribution of specific flavonoid classes, and especially anthocyanins, to resistance against pathogens remains poorly understood.

*Botrytis cinerea* (grey mould) is a devastating necrotrophic plant pathogen with a host range spanning more than 600 plant genera, including vegetables, fruits, ornamentals, and cut flowers (Elad et al., 2016; Singh et al., 2024). This fungus is notorious for its ability to infect aerial tissues such as flowers and leaves, where water-soaked lesions expand into necrotic zones, leading to serious pre-and postharvest losses. While *B. cinerea* is best studied in fruit crops, it also threatens high-value ornamental species. *B. cinerea* infection can induce localised accumulation of flavonoids and anthocyanins. In grape and strawberry, infection has been linked to the localised induction of flavonoid and anthocyanin biosynthesis. For example, in grapes, the “noble rot” that is caused by *B. cinerea* can trigger pink or rosy hues in white grape berries, linked to infection-associated activation of flavonoid and anthocyanin biosynthesis during botrytisation (Blanco-Ulate et al., 2015). Transgenic tomato fruits that were enriched for anthocyanins displayed reduced lesion size and disease severity, while anthocyanin-deficiency resulted in greater susceptibility (Zhang, Butelli, De Stefano, et al., 2013). Nevertheless, the dynamics and functions of flavonoids in floral tissues remain underexplored.

Chrysanthemum (*Chrysanthemum* x *morifolium* Ramat.) is the second most economically important cut flower after roses and is highly susceptible to *B. cinerea*. Grey mould outbreaks drastically reduce marketability by compromising flower quality, appearance, and vase life. Hence, it is important to understand resistance mechanisms against *B. cinerea.* Although chrysanthemum cultivars are known to produce a diverse repertoire of flavonoids and anthocyanins (Lin & Harnly, 2010), their defensive roles against *B. cinerea* have received little attention. Research has been hampered by the polyploid genome of cultivated chrysanthemum. Recent advances in genomic resources now make such investigations feasible. The availability of the diploid *Chrysanthemum seticuspe* genome, alongside a draft assembly of *C. morifolium*, provides a foundation for systems-level analyses of defence responses (Song et al., 2023). Integrating transcriptomic and metabolomic approaches allows for the identification of infection-induced genes and metabolic pathways, as well as the chemical compounds that accumulate in response to pathogen challenge.

Here, we investigated how *C. seticuspe* flowers and leaves respond to *B. cinerea* infection. During evaluation of *C. seticuspe* responses to *B. cinerea*, we noted a striking localised red pigment in inoculated flower petals, seemingly restricting the fungal infection. This observation suggested the possible activation of flavonoid biosynthesis, particularly anthocyanins at pathogen entry sites. We hypothesised that these pigments contribute to floral defence against grey mould. To investigate this, we applied an integrated transcriptomic and metabolomic approach to compare mock-inoculated vs *Botrytis*-inoculated tissues across multiple time points. This study identifies pathogen-induced biosynthetic pathways, characterizes flavonoid and anthocyanin accumulation, and evaluates the antifungal activity of selected metabolites against *B. cinerea*. Our findings provide new insights into the chemical defence strategies of chrysanthemum flowers against grey mould, highlighting floral tissues as active sites of defence and positioning anthocyanins not only as pigments but as key players in defence against *B. cinerea*.

## Materials and methods

### Plant material

The *Chrysanthemum seticuspe* cuttings were obtained from the breeding company Deliflor Chrysanten, Maasdijk, The Netherlands. The cuttings were dipped in rooting hormone IBA and placed in a potting soil under high humidity for 2 weeks. The rooted cuttings were transplanted to potting soil in the greenhouse with temperatures (21°C day; 19°C night) and a relative humidity of 60%.

### *B. cinerea* culture and plant inoculation

*Botrytis cinerea* strain B05.10 was cultured on malt extract agar (MEA) at 20°C for 10 days under a 16 h light/8 h dark photoperiod. Twenty mL of sterile Milli-Q (MQ) water was added to each plate, and conidia were scraped from the surface using a sterile spatula. The resulting mycelial and conidial suspension was filtered through glass wool and transferred to a 50-mL tube. The suspension was centrifuged for 10 min at 1000g to wash the conidia. After discarding the supernatant, the conidia were re-suspended in 30–40 mL of sterile MQ water. Conidial density was determined using a haemocytometer and adjusted to 1 × 10⁷ spores mL⁻¹. Four-week-old leaves and fully opened flowers of *C. seticuspe* were used for *B. cinerea* inoculations. Due to differences in tissue size, leaves were inoculated with 2 µL drops containing 1000 spores µL⁻¹, whereas petals (ray florets) were inoculated with 2 µL drops containing 100 spores µL⁻¹. The inoculation medium consisted of Gamborg B5 (Duchefa, The Netherlands), containing 3.2 g L⁻¹ minerals and vitamins (Gamborg et al., 1976), supplemented with 10 mM sucrose and 10 mM potassium phosphate (pH 6.0). Time-course of petal infection was monitored for red light emission on a Chemi-Doc imager as described in Landeo Villanueva et al., 2021.

### Confocal microscopy

A Leica DMI-8 system paired with a Stellaris-5 confocal laser scanning microscope was used to visualise the localisation of pigmented in relation to fungal infection structures. Petals were inoculated with a GFP-tagged strain of *B. cinerea* and observed at 1 day post inoculation (dpi). Using a white-light laser, samples were excited at 561 nm and emission spectra were recorded between 600-650 nm, which is a suitable range for detection of anthocyanins without including chlorophyll auto-fluorescence signals (Kallam et al., 2017).

### Tissue staining

Staining of inoculated petals at 1 dpi with p-dimethylamino-cinnamaldehyde (DMACA) was done as described by Li et al.,1996. A freshly cut slice of apple was used as positive control. This allowed us to visualise the potential presence of proanthocyanidins, also known as condensed tannins. Staining of fungal hyphae on infected petals was done using aniline blue (Sigma-Aldrich, 2.5% in 2% acetic acid). Samples were spot stained and incubated for 1 hour before washing with MQ. All samples were imaged using a Nikon instrument with D5-5Mc-U2 imaging software at 10x and 40x magnifications with exposure set to ME1/2 s (-1.0 EV).

### Sample preparation for paired omics analysis

Samples of inoculated petals and leaves were taken at different timepoints using 5 mm disposable biopsy punches (Robbins Instruments) to isolate a disc of tissue around the site of inoculation. The collected discs were immediately frozen in liquid nitrogen and stored at-20°C. Three biological replicates were collected per timepoint, with each replicate consisting of at least 10 flower heads or 4–5 fully expanded leaves (6 discs per leaf). The collected samples were freeze-dried and ground to a fine powder in a Retsch MM 400 mixer mill for 5 minutes at 30 Hz using frozen Retsch blocks. For each biological sample, 10 mg of powder was weighed out and distributed into two separate tubes (5 mg each) for transcriptomics and metabolomics analysis, respectively. To monitor instrument stability and technical reproducibility, quality control (QC) samples were also prepared. A composite QC pool was generated by combining powder from randomly selected samples, which was then divided into five separate aliquots. QC samples were processed and analysed in parallel with the biological samples. To extract the metabolites, particularly polar compounds such as anthocyanins, a polar solvent consisting of 75% methanol and 25% dH₂O with 0.1% formic acid was used. A 150 μL volume of extraction solvent was added to each tube. Tubes were vortexed and sonicated for 15 minutes before centrifugation at maximum speed. A 100 μL volume of the supernatant was transferred to glass vials and sealed for mass spectrometry analysis.

### LC-MS/MS analysis

LC-MS/MS was conducted using a Vanquish UHPLC with Exploris120 Orbitrap system (Thermo Scientific), comprising with an Vanquish photodiode array detector (220–600 nm) connected to an Orbitrap Exploris 120 mass spectrometer equipped with an electrospray ionization (ESI) source. The injection volume was 10 μl. Chromatographic separation was performed on a reversed-phase column (Acquity UPLC BEH C18, 1.7 μm, 2.1 × 150 mm; Waters) at 40°C. Degassed eluent A [ultra-pure water: formic acid (1000:1, v/v)] and eluent B [acetonitrile: formic acid (1000:1, v/v)] were used at a flow rate of 0.4 mL/min. A linear gradient of 5 to 75% acetonitrile (v/v) over 22 minutes was applied, followed by 8 minutes of washing and equilibration. FTMS full scans (m/z 90.00–1350.00) were recorded at a resolution of 60,000 FWHM. All the datasets were obtained with positive ionization only.

### LC-MS/MS data analysis

Raw Thermo (.raw) files were converted to mzXML using MSConvert (ProteoWizard v3.0.24207; Chambers et al., 2012). Feature extraction was performed in mzmine v4.4.3 (Schmid et al., 2023) using the ADAP workflow for chromatogram building, deconvolution, isotope grouping, and alignment. (1) Mass detection = MS1 noise level 5.00, MS2 noise level 2.50; (2) ADAP chromatogram builder = minimum consecutive scans 5, minimum intensity for consecutive scans 4.0E5, minimum absolute height 2.0E6, m/z tolerance scan-to-scan 0.0020 m/z or 10.0 ppm; (3) Smoothing = Savitzky Golay; (4) Local minimum feature resolver = chromatographic threshold 90.0%, Minimum search range RT/Mobility 0.040 min, minimum absolute height 2.0E6, min ratio of peak top/edge 2.00, peak duration range 0.00–1.20 min, minimum scans 5; (5) 13C isotope filter = m/z tolerance 0.0015 m/z or 3.0 ppm, retention time tolerance 0.04 min, monotonic shape yes, maximum charge 2, representative isotope most intense, never remove feature with MS2 on; (6) Isotopic peaks finder = m/z tolerance 0.0015 m/z or 3.0 ppm, maximum charge of isotope m/z 1, search in scans single most intense; (7) Join aligner = m/z tolerance 0.0015 m/z or 5.0 ppm, retention time tolerance 0.10 min; (8) Feature list rows filter = Minimum aligned features (samples) max of 2 samples or 50.0%; (9) Peak finder (gap filling) = intensity tolerance 20.0%, m/z tolerance 0.0020 m/z or 10.0 ppm, retention time tolerance 0.10 min, minimum scans 3 (mzmine batch file is available on Zenodo - see Data availability section). Features detected from mzmine were used for downstream analysis, and the resulting.mgf file and feature quantification table were exported to GNPS2. A Feature-Based Molecular Network (FBMN) was performed on GNPS2 (gnps2.org) (Nothias et al., 2020), the successor platform of GNPS (Wang et al., 2016), using default parameters mentioned below: precursor ion mass tolerance of 2.0 Da, fragment ion mass tolerance 0.5 Da, minimum cosine score that must occur between a pair of MS/MS spectra in order to form an edge in the molecular network of 0.7, minimum number of fragment ions that are shared between pairs of related MS/MS spectra in order to be connected by an edge in the molecular network of 6, maximum shift between precursors 1999 Da, maximum number of neighbour nodes for one single node of 10, maximum size of nodes allowed in a single connected network 100; search analogs op, minimum number of fragments that MS/MS spectra should contain in order to be considered for annotation of 6, score threshold of 0.7. The network was visualized in Cytoscape 3.10.1 (Shannon et al., 2003), and the job is accessible at https://gnps2.org/status?task=91fe83a1f1ca41f1acb7d5512476e65c. Additionally, all MS/MS-containing features were annotated using SIRIUS v6.1 (Dührkop et al., 2019), all possible adducts. The SIRIUS workflow included ZODIAC for molecular-formula identification, CANOPUS for compound-class prediction (Dührkop et al., 2021), NPClassifier for natural-product class assignment (Kim et al., 2021), CSI:FingerID (Dührkop et al., 2015) all fall back adducts, structure-similarity searches against the BioCyc, PubChem, HMDB, COCONUT, LOUTS, ChEBI, KEGG databases.

### Statistical analysis

Quantitative feature tables were processed using FBMNstats (Pakkir Shah et al., 2025). Blank subtraction and random-value imputation were applied prior to log₂ transformation. Features were mean-centred across samples. Differential accumulation was tested using the Kruskal–Wallis test followed by Dunn’s post hoc test (p < 0.05).

### Transcriptomic analysis

Total RNA was extracted using a Maxwell 16 LEV Plant RNA Kit (Promega, USA). RNA sequencing was performed at the Beijing Genomics Institute (BGI, Shenzhen, China). Raw data and metadata is publicly available under NCBI BioProject accession number PRJNA1369786. Clean reads were mapped to the *Chrysanthemum seticuspe* ‘Gojo-0’ genome (Nakano et al., 2021) using HISAT2 (Kim et al., 2019). Transcript assembly and quantification were conducted with StringTie (Pertea et al., 2015), and read counts were used for differential expression analysis with DESeq2 (Love et al., 2014). Genes showing |log₂(fold change)| > 2 and p < 0.01 were considered significantly differentially expressed. Biosynthetic gene clusters (BGCs) were identified using PlantiSMASH 1.0 version (Kautsar et al., 2017) with default parameters. Heatmaps were constructed from TPM values and expression data were row-scaled (z-scores) across all samples. KEGG annotation for the *C. seticuspe* proteome (CsGojo, obtained from MumGARDEN (https://mum-garden.kazusa.or.jp) was obtained using BlastKOALA (Kanehisa et al., 2016), using the eukaryotic reference gene database, filtering for pathways present in plants. Enrichment for KEGG pathways in differentially expressed genes was tested using Fisher’s exact test (implemented in SciPy (Virtanen et al., 2020)). P-values were corrected for multiple testing using Benjamini-Hochberg False Discovery Rate correction (implemented in StatsModels (Seabold & Perktold, 2010)).

### Functional annotation and identification of genes in *C. seticuspe*

Flavonoid biosynthetic genes in *C. seticuspe* were predicted using KIPEs (Rempel et al., 2023) with default parameters. Candidate genes were selected based on the highest similarity scores, with priority given to conserved amino acid residues and functional domains. When multiple candidates for a given enzyme did not share fully conserved residues, the top-scoring sequence was retained for further analysis. Transcription factors were identified using bHLH annotator (Thoben & Pucker, 2023) and MYB annotator (Pucker, 2022).

### Resazurin-based assays for antifungal activity of flavonoid and anthocyanin compounds

The antifungal activity of commercially available flavonoid and anthocyanin compounds was tested using an assay based on the conversion of non-fluorescent resazurin dye (Sigma R7017) to fluorescent resorufin by metabolic activity of *B. cinerea*. In a 96-well plate, the fungal spore suspension was combined with RPMI medium and the antifungal compound at the desired concentration. Fluorescence emission was measured at 615 nm after excitation at 570 nm. Data was collected in CLARIOStar MARS software and processed in R. Fungal growth was normalised based on the maximal fluorescence obtained without the antifungal compound (concentration 0 μg/mL) and averaged per concentration. Standards for the following flavonoids and anthocyanins were purchased from Sigma-Aldrich: Tilianin (PHL85779), Acacetin (00017), Quercetin (337951), Hyperoside (PHL89227), Cyanidin chloride (79457), Cyanidin 3,5-diglucoside chloride (PHL89615).

### Multi-omics analysis using MEANtools

Upon normalization of transcriptomic and metabolomic matrices, the MEANtools (Singh et al., 2025) (version 0.1.0) (https://github.com/kumarsaurabh20/meantools) workflow was applied to generate transcript-metabolite networks and predict biosynthetic pathway steps. First, mass features from the metabolic feature file were queried against the LOTUS (Rutz et al., 2022) database using the *queryMassNPDB.py* script, assigning tentative structural annotations to each m/z value considering most common adduct formations. To refine pathway predictions based on taxonomic relevance, we ran the MEANtools workflow twice using different taxonomy-based filter settings. In the first run, mass features were queried, and reaction-rule mappings were filtered using “Chrysanthemum” as the species-level criterion; in the second run, a broader taxonomic category “Asteraceae” was used to capture reactions conserved across the family. All other parameters remained at their default settings. Next, the *corrMultiomics.py* script calculated Pearson correlations between transcript expression and metabolite abundances and converted these into Mutual Rank (MR) scores, which were then transformed into continuous edge-weights across multiple decay rates (e.g., DR = 5, 10, 25, 50). Functional clusters (FCs) were identified in each network via the transcript-metabolite clustering algorithm implemented in MEANtools; FCs were manually merged using based on shared metabolites and available annotations between FCs, producing a refined set of transcript-metabolite pairs for downstream analysis. The *pathMassTransitions.py* script mapped filtered mass transitions from the RetroRules database (RR) (Duigou et al., 2019) and MetaNetX (Moretti et al., 2021) against the observed metabolome, creating substrate-product pairs consistent with the dataset. The *heraldPathways.py* script predicted reaction steps across multiple iterations. For each taxonomy-specific dataset, pathway prediction was performed with two iteration depths to explore pathway continuity: once with *i = 3* and again with *i = 5*, enabling comparison of short-range versus extended biosynthetic trajectories. These complementary analyses allowed us to assess the stability of predicted reaction routes and evaluate how taxonomic scope and pathway iteration depth influence the reconstruction of putative biosynthetic pathways from paired transcriptomics-metabolomics data. Finally, *paveWays.py* script generated schematic visualizations and reaction-likelihood scores to prioritize plausible enzymatic steps, thereby enabling translation of multi-omics correlations into mechanistic pathway. It produced SVG schematic files and tables allowing us to query specific metabolite sets and interpret the structural relationships within predicted routes.

## Data availability

Raw data from the RNA-seq is publicly available under NCBI BioProject accession number PRJNA1369786. LC–MS/MS files (positive mode) were deposited in MassIVE under accession number MSV000100208. The mzmine batch file, processed feature tables, annotation tables, SIRIUS outputs, MEANtools output, metadata have been deposited in Zenodo: 10.5281/zenodo.17910126

## Results

### Localised red pigmentation coincides with fungal penetration sites in flower petals

After inoculation of *B. cinerea* on flower petals, localised red spots appeared as early as 1-day post-inoculation (dpi), directly beneath the inoculation droplets. The number of red spots significantly increased as the infection proceeded to 3 dpi (Figure 1A). At higher magnification, each spot corresponded to a discrete cluster of red-pigmented epidermal cells surrounding a fungal penetration site. Aniline blue staining confirmed the presence of *B. cinerea* hyphae above the pigmented cells. Confocal microscopy of petals inoculated with *B. cinerea* expressing cytoplasmic GFP further revealed that infection structures (infection cushions and penetration pegs) co-localised with intense host autofluorescence in the red channel (561 nm excitation; 600–650 nm emission) (Figure 1B). In contrast, inoculated *C. seticuspe* leaves did not develop red pigmentation, but instead formed expanding water-soaked necrotic lesions by 3 dpi (Figure 1C). Time-course red-light imaging on a ChemiDoc MP (green LED excitation; 605/650 nm emission filter) detected emission at petal inoculation sites from 1 dpi, which intensified by 3 dpi, consistent with progressive fungal colonization and host cell death. Since anthocyanins are known to fluoresce in this spectral range (Kallam et al., 2017), the red-channel signal may reflect localised anthocyanin accumulation, a possibility that was further evaluated by LC-MS/MS analysis.

**Figure 1:**
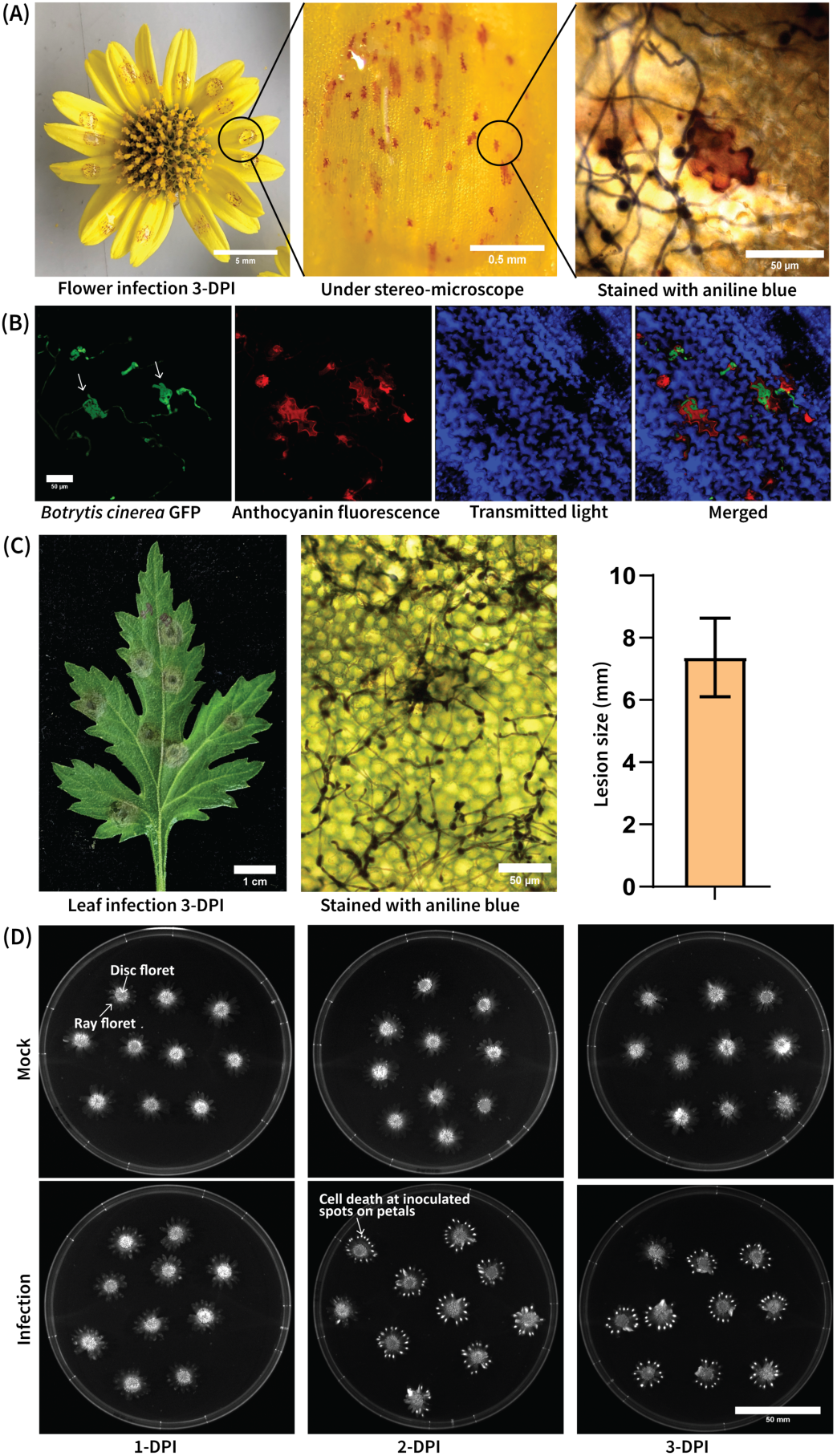
Colonisation and lesion development of *Botrytis cinerea* on *Chrysanthemum seticuspe* flower petals and leaves. (A) Infection of *C. seticuspe* flower petals by *B. cinerea* at

### Infection triggers transcriptional reprogramming of specialised metabolism

To investigate the underlying molecular responses accompanying these symptoms, we performed transcriptomic profiling of mock-trated and *B. cinerea* inoculated tissues of chrysanthemum flowers and leaves. Principal component analysis (PCA) of RNA-seq expression profiles showed a clear separation between infected and control samples, especially in flower petals at 24, 48, and 72 hours post-inoculation (hpi) (Figure 2A). A similar distinct clustering was observed in leaves at 12 and 20 hpi (Figure 2B), indicating that *B. cinerea* infection elicited substantial changes in gene expression in both organs. Thousands of genes were differentially expressed upon infection, with many upregulated genes associated with stress and defence responses. Notably, KEGG pathway enrichment analysis revealed that multiple secondary metabolism pathway genes were significantly up-regulated in infected tissue. Particularly the sesquiterpenoid, phenylpropanoid and flavonoid biosynthetic pathways were significantly induced in infected tissues compared to controls (Figure 2C and 2D). In addition, a plantiSMASH analysis of the *C. seticuspe* genome identified several putative secondary metabolite biosynthetic gene clusters (BGCs); many of the clusters showed coordinated elevated expression specifically in infected petals (Figure 2D). Numerous terpene, alkaloid and polyketide gene clusters were upregulated in infected flowers as early as 24 hpi suggesting activation of dedicated metabolic pathways during the infection in flowers.

**Figure 2:**
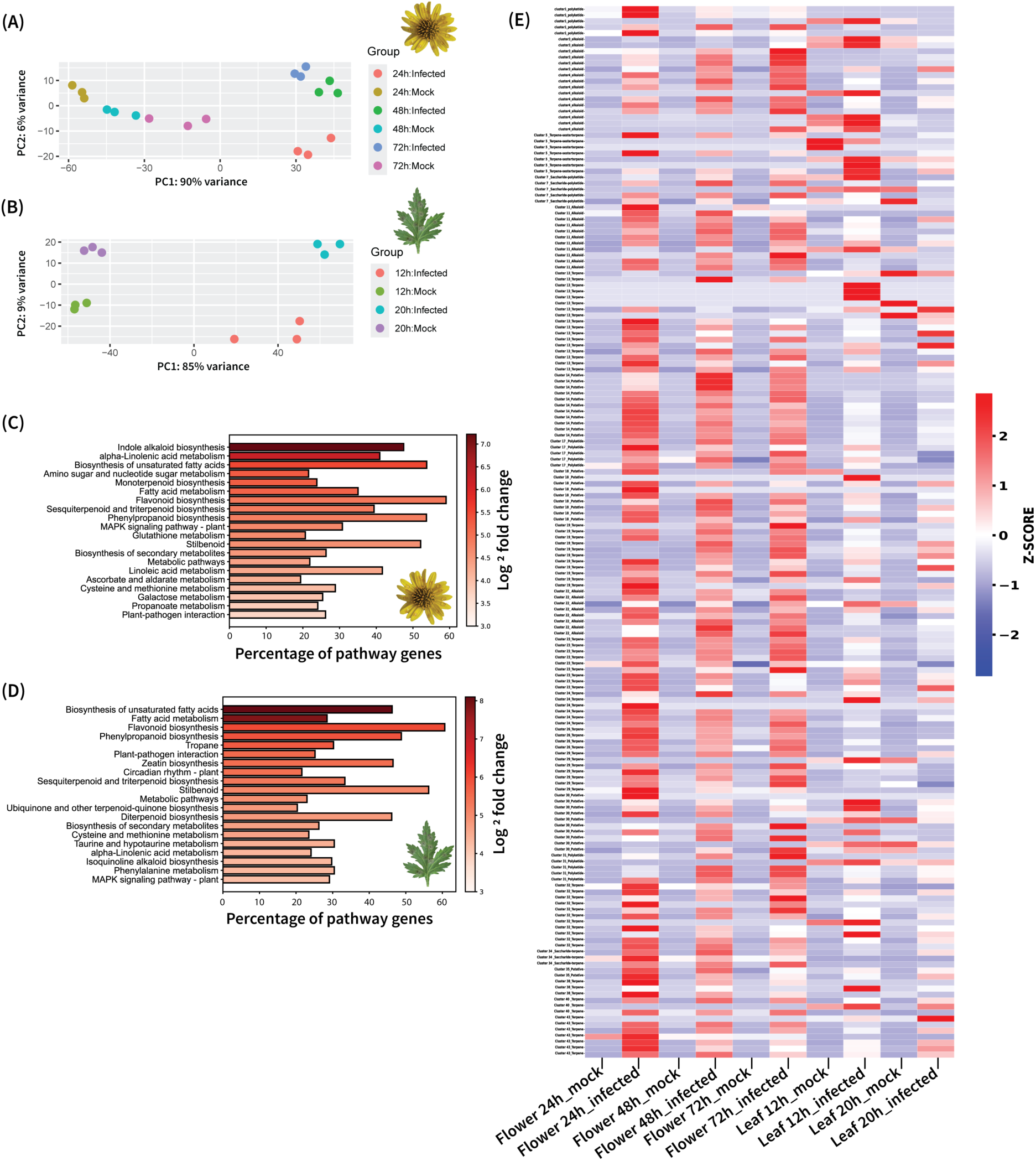
Transcriptomic responses of Chrysanthemum seticuspe petals and leaves to Botrytis cinerea infection. **(A)** Principal component analysis (PCA) of RNA-seq profiles from petals (Mock vs *B. cinerea* infected) at 24, 48, and 72 hours post-inoculation (hpi). **(B)** PCA of RNA-seq profiles from leaves (Mock vs *B. cinerea* infected) at 12 and 20 hpi. Individual dots represent sample replicates. **(C and D)** KEGG pathway enrichment of differentially expressed genes (DEGs) in *B. cinerea* infected petals and leaves relative to mock controls. Bars indicate the number of DEGs assigned to each pathway; colour intensity reflects the average log₂ fold-change of DEGs within each pathway (FDR < 0.05). **(E)** Heatmap of putative BGCs predicted by plantiSMASH. Expression values are row-scaled z-scores across all samples (petals: 24, 48, 72 hpi; leaves:12, 20 hpi).

### Flavonoid and anthocyanin pathway genes are selectively induced in infected petals

One of the most prominent infection-responsive pathways was the flavonoid/anthocyanin biosynthetic pathway. Consistent with the visible red pigmentation, multiple genes encoding anthocyanin biosynthetic enzymes together with putative MYB–bHLH–WD40 (MBW) regulatory genes, were strongly upregulated in infected petals. A heatmap of gene expression (Figure 3C) showed that upon *B. cinerea* inoculation, transcripts for most key enzymes accumulated to higher levels in petals but not in leaves, in line with the absence of anthocyanin pigmentation in leaves. Several candidate MBW transcription factors, including R2R3-MYB and bHLH genes, displayed infection-induced expression in petals, alongside anthocyanin structural biosynthetic genes. Although KIPEs (Rempel et al., 2023) predicted two anthocyanidin synthase (ANS) genes, only one possessed all conserved residues. However, neither of these genes showed detectable expression under any condition, suggesting that an alternative set of genes might fulfil this role. Notably, the *C. seticuspe* orthologs of Arabidopsis bHLH transcription factors TT8 and MYC1, both canonical components of the MBW complex, showed no induction after infection either, suggesting the possible involvement of an alternative set of transcription factors in regulating anthocyanin accumulation during *C. seticuspe* petal infection.

**Figure 3:**
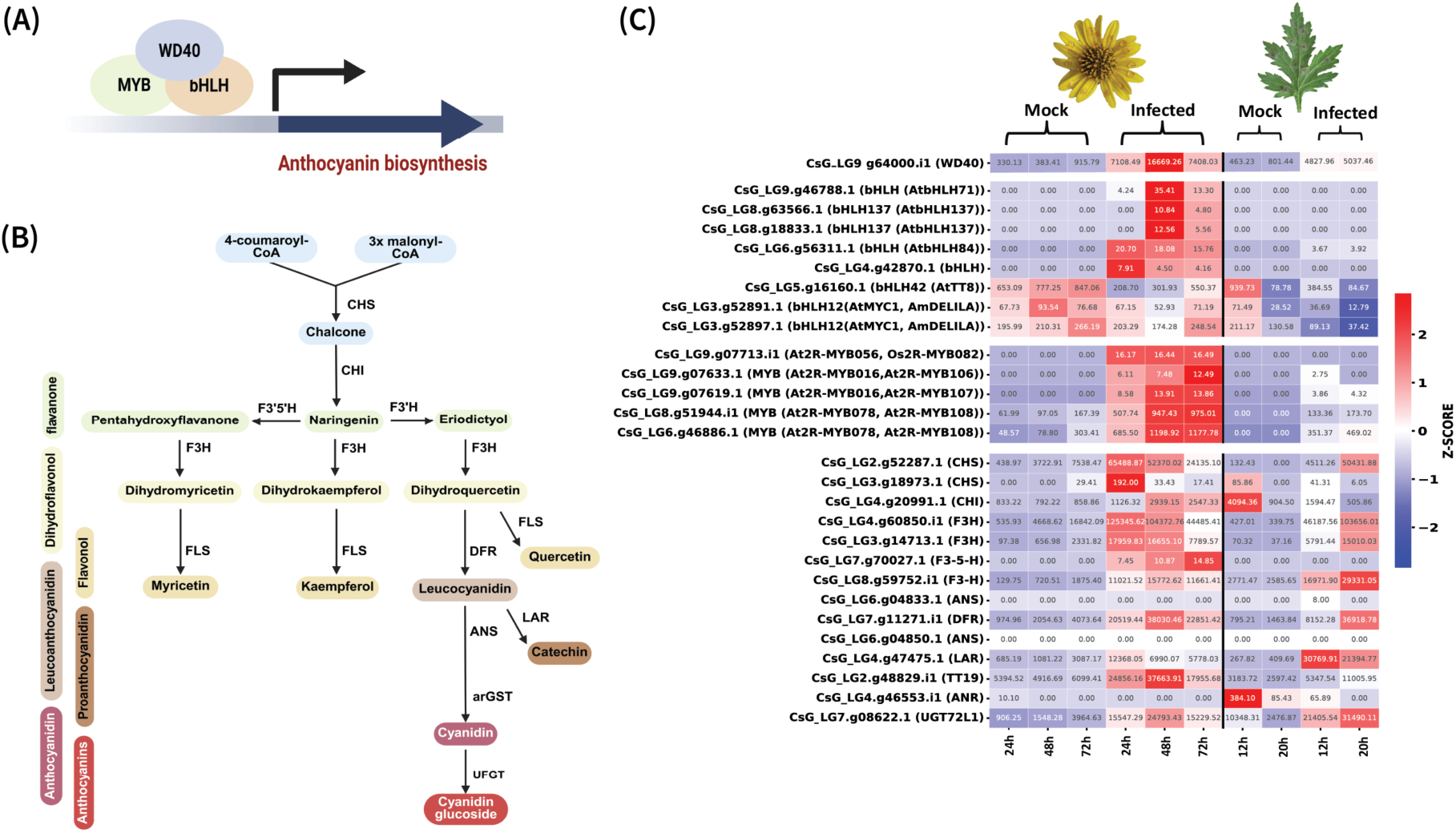
Pathogen-induced activation of the anthocyanin biosynthetic pathway in *Chrysanthemum seticuspe*. **(A)** Model of MYB–bHLH–WD40 (MBW) transcriptional complex regulating anthocyanin biosynthesis in *Arabidopsis* (and other model plants) through binding to promoters of structural genes. **(B)** Simplified anthocyanin biosynthetic pathway. Starting from 4-coumaroyl-CoA and three molecules of malonyl-CoA, chalcone synthase (CHS) and chalcone isomerase (CHI) generate naringenin, which is hydroxylated by flavanone 3-hydroxylase (F3H) into dihydroflavonols. Subsequent reactions catalyzed by dihydroflavonol 4-reductase (DFR) and anthocyanidin synthase (ANS) yield anthocyanidins, which are stabilized into anthocyanin glucosides by flavonoid 3-O-glucosyltransferase (UFGT) using UDP-glucose. Side branches of the pathway lead to flavonols (via flavonol synthase, FLS) and catechins (via leucoanthocyanidin reductase (LAR) and anthocyanin-related glutathione S-transferase (arGST). **(C)** Heatmap of z-score normalised expression for candidate MBW regulators and anthocyanin structural genes (predicted by KIPES). Values inside the box are TPM values. Rows indicate gene identifiers with putative annotation in parentheses;columns show samples from petals (mock vs *B. cinerea* infected at 24, 48, 72 hpi) and leaves (mock vs infected at 12, 20 hpi).

### Metabolite profiling highlights petal-specific accumulation of flavonoids and terpenoids

Metabolomic profiling of infected plants mirrored the transcriptomic changes, revealing a pronounced accumulation of specialised metabolites in petals. PCA of untargeted metabolite profiles showed that infected petal samples clustered distinctly from mock-inoculated controls at all timepoints, reflecting pronounced infection-induced metabolic shifts. In contrast, infected leaves showed less separation from controls in the PCA, and the overall magnitude of changes was less pronounced (Figure 4A and supplementary Figure S3). Among the many metabolite features detected, those that changed significantly upon infection were predominantly annotated as belonging to the terpenoid, fatty acid, and shikimates-phenylpropanoids classes (Figure 4B). In petals, a substantial subset of differentially abundant metabolites were the flavonoids (Figure 4C), a finding consistent with transcriptional activation of the flavonoid and anthocyanin biosynthetic pathways described above. By contrast, infected leaves exhibited only minimal induction of these metabolite classes. Together, the data indicate that *B. cinerea* infection triggers strong reconfiguration of the petal metabolome, characterized particularly by the accumulation of terpenoids and flavonoids, whereas the metabolic response in leaves was comparatively limited.

**Figure 4:**
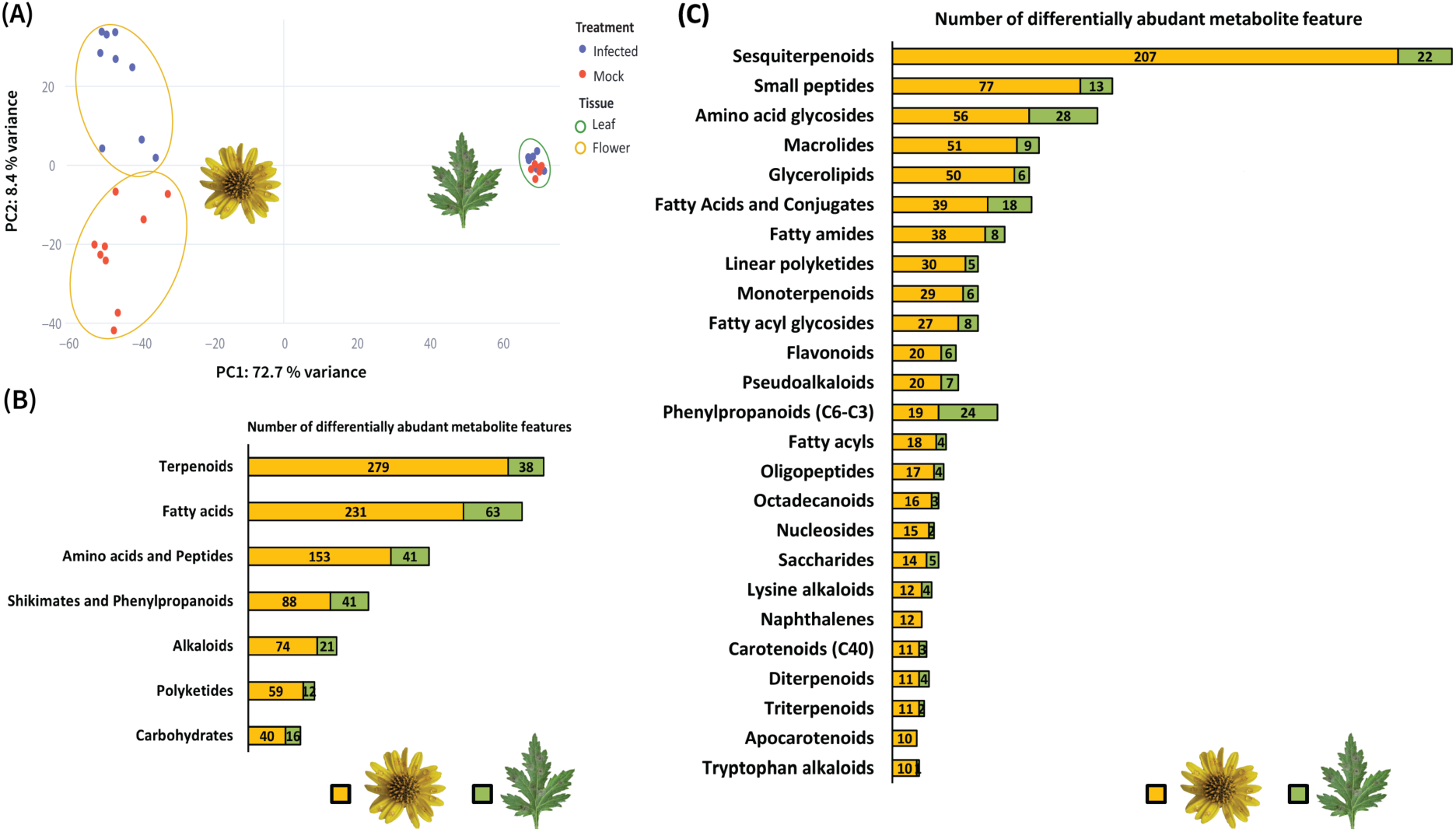
Metabolome remodelling in Chrysanthemum petals vs leaves upon *B. cinerea* infection. **(A)** Principal component analysis (PCA) of metabolite features from mock-inoculated (red) vs *B. cinerea*-infected (blue) samples. Petal samples are outlined in gold (24, 48 and 72 h post-inoculation, hpi) and leaf samples in green (12 and 20 hpi). **(B)** Comparison of metabolite super-classes (NPClassifier) differentially abundant upon infection, shown for petals (left) and leaves (right). **(C)** Distribution of differentially abundant metabolite features of top 25 subclasses (NPClassifier). Bars show the counts of infection-responsive features for petals (left) and leaves (right). Differential abundance was determined using the Kruskal–Wallis test followed by Dunn’s post-hoc test (*p* < 0.05). For tissue-specific PCA plots and pooled quality control (QC) assessment, see Supplementary Figure S3.

### Time-resolved profiling reveals coordinated induction of defence-related flavonoids and anthocyanins

Consistent with the global metabolomic shifts, specific flavonoid and anthocyanin compounds accumulated to high levels in infected petals over the course of infection. Several metabolites that were barely detectable in mock-treated petals showed strong time-dependent induction in infected petals (Figure 5). For example, the flavonol aglycone quercetin increased sharply in concentration by 48–72 hpi in infected petals, while it remained low in controls (Figure 5A). Its 3-O-glycosylated derivative hyperoside (quercetin 3-galactoside) was similarly elevated in infected tissues (Figure 5D). The flavone tilianin and its aglycone acacetin both showed progressive accumulation in petals following inoculation (Figure 5B, 5E). Notably, an anthocyanin candidate identified by SIRIUS as cyanidin 3-glucoside-5-(6-acetylglucoside), accumulated to high levels in infected petals by 72 hpi, but in mock inoculated petals remained at baseline (Figure 5C), which corresponds with the visual increase of red spots at 72 hpi. Together, the coordinated temporal rise of diverse flavonoids and anthocyanins highlights a strong petal-specific metabolic response mounted by *C. seticuspe* against *B. cinerea*.

**Figure 5:**
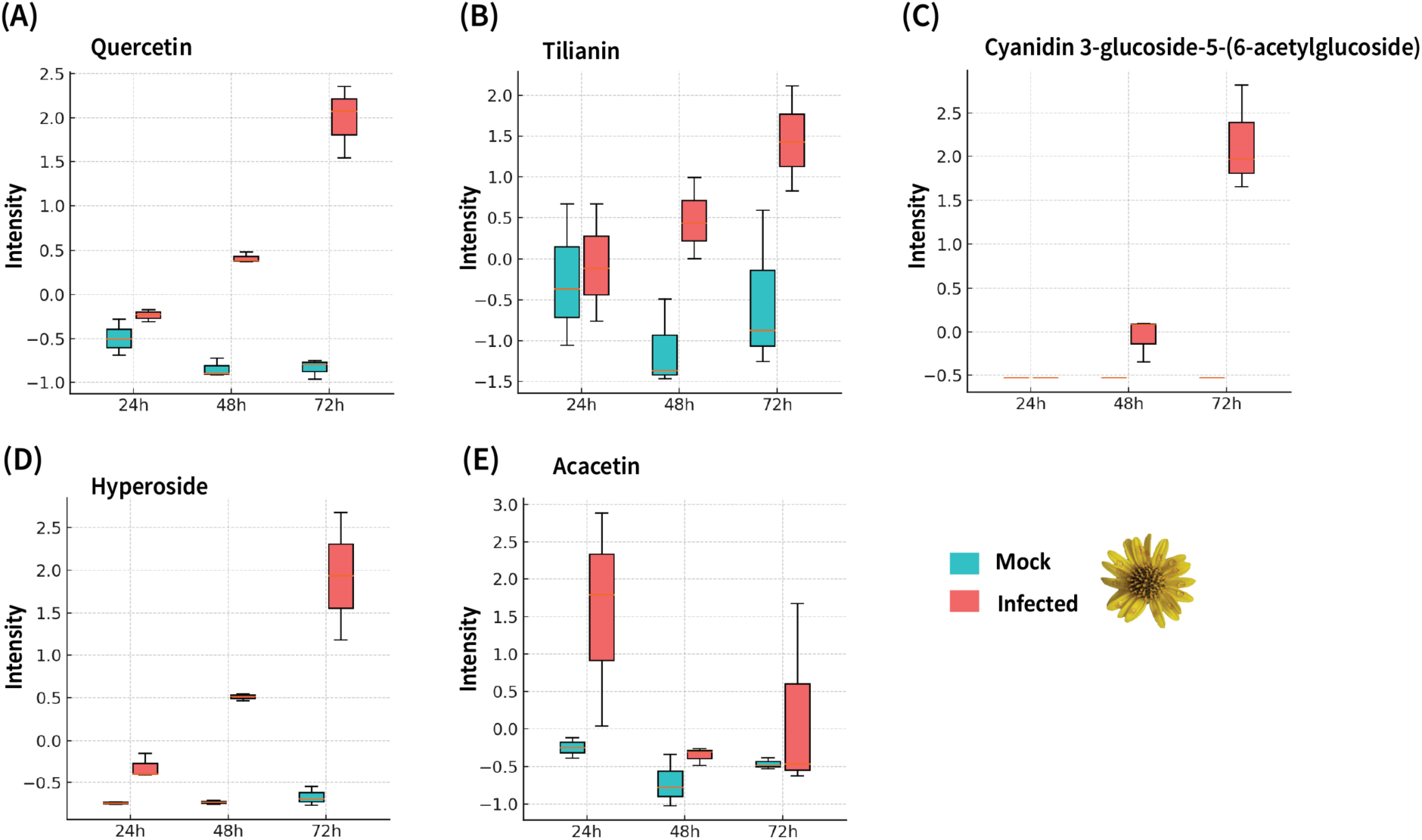
Time-course of accumulation of diSerentially up-regulated flavonoids and anthocyanins from the untargeted metabolomics of Chrysanthemum petals following *B. cinerea* infection. Boxplots show z-score normalized intensities of five metabolites significantly induced in infected petals (red) compared to mock controls (teal) at 24, 48, and 72-hours post-inoculation (hpi). (A) Quercetin, (B) Tilianin, (C) Cyanidin 3-glucoside-5-(6-acetylglucoside), (D) Hyperoside (hyperin) and (E) Acacetin. Each box represents the interquartile range (IQR) with the median line; whiskers extend to 1.5× IQR. Infected petals show progressive accumulation of all five metabolites across the time course. Differential accumulation was tested using the Kruskal–Wallis test followed by Dunn’s post hoc test (p < 0.05).

### Pathogen-induced flavonoids and anthocyanins exhibit antifungal activity

To determine whether the flavonoids and anthocyanins that accumulate during infection can directly inhibit *B. cinerea*, we tested commercially available pure compounds for several abundant metabolites detected in infected petals. The tested compounds included the aglycones of two flavonoids (quercetin and tilianin), as well as an anthocyanin (cyanidin) and its glycosylated form. We were unable to purchase the infection-specific compound cyanidin 3-glucoside-5-(6-acetylglucoside). Instead, we tested the cyanidin disaccharide conjugate cyanidin 3,5-diglucoside. Fungal growth inhibition assays showed that each of the tested compounds inhibited *B. cinerea* growth in a dose-dependent manner (Figure 6), confirming that both flavonoids and anthocyanins possess antifungal activity. The flavonoids quercetin and acacetin inhibited *B. cinerea* growth to a similar extent as their respective glycosides, hyperoside and tilianin (Figure 6A, 6B). In contrast, the aglycone cyanidin displayed significantly higher antifungal potency than its disaccharide conjugate, cyanidin 3,5-diglucoside (Figure 6C). Collectively, these results demonstrate that both the aglycones and their glycosylated derivatives possess direct antifungal activity, contributing to the chemical defence against grey mould.

**Figure 6:**
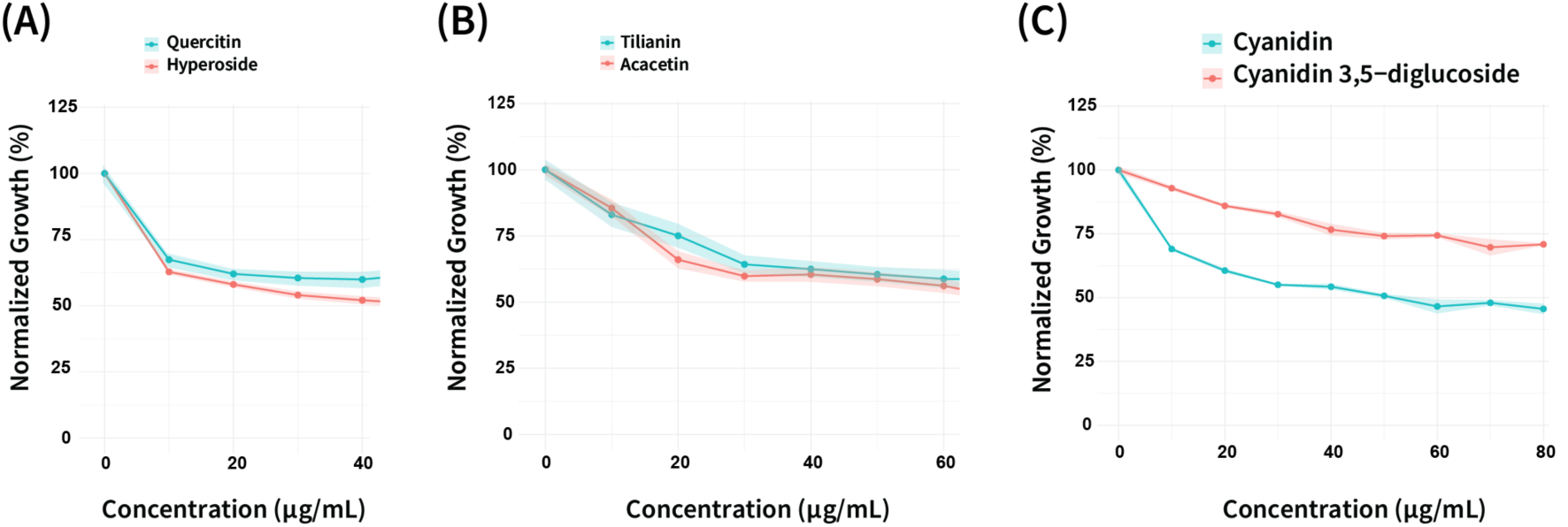
Concentration-dependent inhibition of *B. cinerea* growth by flavonoids and anthocyanins, measured via resazurin assay. Fungal biomass (mean ± SD, n = 3) was quantified after 20h incubation at increasing metabolite concentrations. Values are normalised to untreated controls (100%). (A) Quercetin (teal) vs Hyperoside (red). (B) Tilianin (teal) vs Acacetin (red). (C) Cyanidin (teal) vs Cyanidin 3,5-diglucoside (red).

### Transcript-metabolite co-clustering with MEANtools reveal new biosynthetic pathways

Paired omics datasets are well suited to study biosynthetic routes through pathway discovery linking genes and molecules. Here, we used our time-course RNA-seq and LC–MS/MS data as input for MEANtools (Singh et al., 2025), which builds a joint network that links metabolite abundance to gene expression and proposes biosynthetic pathways. The full MEANtools network contained several transcript-metabolite modules or functional clusters. One of the merged clusters was strongly enriched for known anthocyanin pathway genes. We refer to this as the anthocyanin module (Figure 7A) and use it as a case study for how MEANtools can highlight a pathogen-induced biosynthetic pathway. The anthocyanin module network shows a modular topology organised into several biosynthetic and regulatory sub-graphs. The network has a scale-free topology, with a small set of highly connected hub transcripts that link many dense local clusters of co-regulated enzymes and transporters. The node-degree distribution is strongly right-skewed (Supplementary Figure S2A), so most transcripts in the transcript-metabolite network connect to only one or a few metabolites, whereas a minority of genes act as hubs connected to many metabolites. This pattern matches current views of plant metabolic wiring, where pathway enzymes (for example in the anthocyanin or phenylpropanoid routes) tend to have focused connections, and regulatory or transport genes link several modules. The relation between node degree and edge weight (Supplementary Figure S2B) does not follow a simple positive trend. Instead, the scatter plot has a bell-shaped form: genes with many edges tend to show intermediate average association strength, consistent with broad but less specific regulation. This category includes genes encoding global co-regulators or enzymes that act at pathway branch points. Genes with few edges often show very strong or very weak associations. This points to specialised or condition-dependent roles, in which genes are tightly coupled to a small set of metabolites (for example, enzymes catalysing individual reactions), whereas others show weak links that may reflect peripheral or indirect associations.

**Figure 7:**
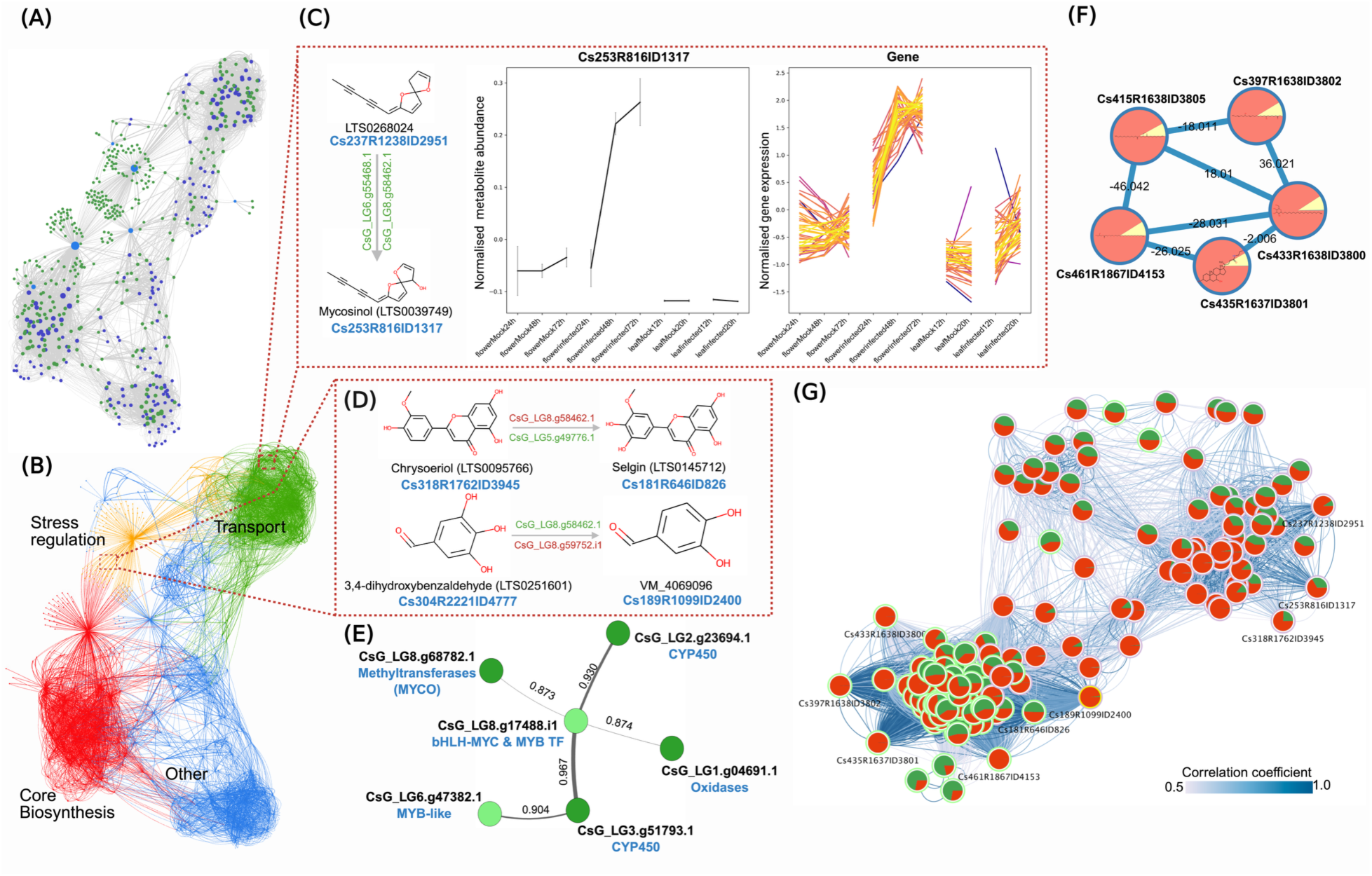
Transcript-metabolite interaction network links an anthocyanin defence module to mycosinol-like metabolites. **(A)** Anthocyanin transcript–metabolite mutual-rank–based network showing co-regulated modules (green, transcripts; blue, metabolites). **(B)** The same network annotated into functional domains representing core biosynthesis (red), regulation of stress response (orange), transport (green) and other associated functions (blue). **(C)** MEANtools-predicted mycosinol metabolite Cs253R816ID1317 and related structure with LOTUS ID (LTS0268024) as source of putative annotation with “Asteraceae” as the filtering step in the LOTUS database. Left, chemical structures with candidate tailoring genes predicted my MEANtools; center, normalised abundance of metabolite Cs253R816ID1317 in mock and infected samples; right, z-score normalised expression of genes correlated with Cs253R816ID1317. **(D)** Additional MEANtools-prioritised biosynthetic hub in the anthocyanin module, including chrysoeriol (LTS0059766), selgin (LTS0145712), 3,4-dihydroxybenzaldehyde (LTS0251601) and VM_4060906 a virtual molecule predicted by MEANtools with “Chrysanthemum” as the filtering parameters in the LOTUS database. **(E)** Local gene subnetwork linked to Cs253R816ID1317. Nodes include methyltransferases, MYB/bHLH transcription factors, cytochrome P450s and oxidases; edge labels give correlation coeoicients. **(F)** GNPS molecular network of sphingolipid candidates (infection-enriched features) correlating with the putative mycosinol-related pathway. Nodes show relative abundance in infected leaf (pink) and infected flowers (yellow); the edges are Δm/z values. **(G)** A network representing high-correlation between metabolites from the anthocyanin module. Nodes represent metabolic features, pie-chart contains abundance of feature in mock (green) vs infected (red) samples. The edges show medium to high correlation (0.5-1.0). Metabolites from mycosinol-like, intermediate steps.

Inside the anthocyanin module, four functional sub-modules can be distinguished (Figure 7B). A core anthocyanin biosynthesis cluster (including CHS, CHI, F3H, DFR, ANS and UFGT homologues) forms the metabolic backbone and connects to glycosyltransferases that diversify pigment structures. A stress-related cluster links flavonoid biosynthesis to wider phenolic metabolism and contains caffeoyl-CoA O-methyltransferases and hydroxycinnamoyl transferases that feed lignin-like branches. A membrane and transport cluster contains ATP-binding cassette (ABC) and MATE transporters, consistent with compartment-specific regulation and vacuolar sequestration of flavonoids. GO and PFAM analysis in these clusters shows strong enrichment of oxidoreductase activity (GO:0016491), UDP-glucosyltransferase activity (GO:0035251) and transport-associated domains (PF00005, PF01554), in line with redox-controlled steps and vacuolar transport in anthocyanin and related flavonoid metabolism. Transcription factor categories enriched in the same region of the network include MYB–bHLH–WD40 complexes, in agreement with the layered control within this metabolic pathway of structural and regulatory gene sets. We combined the transcript-metabolite network with the reactions proposed by MEANtools. Searches based on “Asteraceae” suggested that many of the predicted reactions may act on mycosinol-like metabolites related to polyyne-type phytoalexins (Supplementary material, MEANtools_output in Zenodo). Mycosinol is an anti-fungal (spiroketalenol ether) phytoalexin known to be produced by *Coleostephus myconis* (Asteraceae) after infection with *B. cinerea* (Marshall et al., 1987).

Searches based on “Chrysanthemum” highlighted metabolic features tentatively assigned as chrysoeriol and selgin (both flavones) and a virtual molecule (phenolic aldehyde) predicted by MEANtools. These compounds had strong correlations with sphingolipids, which could represent intermediates or side branches of the mycosinol route, all of which clustered within the GNPS molecular network (Figure 7F). All these metabolites are enriched in infected tissues (flowers and leaves) and map to the stress-related submodule of the anthocyanin network, where they connect to MYB/bHLH transcription factors, methyltransferases and oxidases (Figure 7B,D,E). Together with the presence of flavone-like conjugates, this suggests that flavone conjugates, phenolic aldehydes and sphingolipid-like features participate in a common infection-induced stress response branch, that is transcriptionally linked to the anthocyanin module. Anthocyanins and mycosinol arise from distinct metabolic backgrounds, the phenylpropanoid and fatty-acid pathways, respectively. Even though the structural pathways are different, the integrated transcript-metabolite network often has cross-links between anthocyanins and polyyne modules through shared redox / tailoring enzyme and coexpression/coregulation under stress (Figure 7E). This means that in the integrated network, it is likely for anthocyanin biosynthetic genes to cluster near mycosinol-related oxidases not because they share substrates, but because they share transcriptional triggers.

Metabolites prioritised through MEANtools showed strong metabolite-metabolite associations (Figure 7G) and this association aligned well with one of the GNPS molecular network results (Figure 7F). For example, metabolite 2951 displayed a very strong correlation with metabolite 1317 (r = 0.92, p = 7.11×10⁻¹³, Mutual Rank = 16), and both were putatively annotated as mycosinol-like compounds from the LOTUS database (Figure 7G). These compounds were highly abundant in infected samples compared with mock controls (Figure 7C). Similarly, metabolites 3945 and 826 showed a weak but negative correlation (r = –0.38, p = 0.03), which may reflect a flux-based relationship where an increase in one metabolite corresponds to a decrease in the other. Metabolite 3945 was strongly correlated with 2951 (r = 0.61, p = 3×10⁻⁴) and 1317 (r = 0.54, p = 4.5×10⁻⁴) and showed structural similarity to 826 predicted from the LOTUS database. In summary, the combined topology, enrichment patterns and abundance profiles place the anthocyanin module at the centre of redox-linked defence. MEANtools networks were exported as interactive HTML files for detailed inspection and manual curation, and all predicted reactions are supplied in the supplementary material made available in Zenodo (see Data availability section).

## Discussion

Flower tissues are often considered more susceptible to fungal pathogens than leaves, yet we observed that *Chrysanthemum seticuspe* flower petals mounted an active defence that restricted *B. cinerea* at the infection sites. After inoculation with *B. cinerea*, discrete red spots appeared at the pathogen penetration sites within 24hrs post inoculation and intensified over time. These pigmented spots co-localised with fungal infection structures (penetration pegs and infection cushions) and anthocyanin autofluorescence was observed in epidermal cells. In contrast, infected leaves developed expanding necrotic lesions without visible pigments. Together, these observations indicate an early, spatially confined defence response in the flower petals of *C. seticuspe* that is macroscopically visible at infection sites.

This effective resistance response links to a massive, petal-specific transcriptional reprogramming that is distinct from the leaves. While both organs activate general defence genes, the petal transcriptome converged on the upregulation of several specialised secondary metabolic pathways, particularly those involved in the biosynthesis of terpenoids, phenylpropanoids, flavonoids, and anthocyanins. Several predicted secondary-metabolite gene clusters (mostly terpenoids) showed coordinated induction in the petals. Chrysanthemum flowers are known for their scent, and our metabolomics data showed terpenoids as the largest compound class detected, consistent with reports that chrysanthemum floral volatiles are dominated by terpenoids (He et al., 2023). The features associated with the red phenotype were linked to flavonoids and anthocyanins. In infected petals, flavonoids (quercetin, hyperoside, acacetin and tilianin) and the anthocyanin cyanidin 3-glucoside-5-(6-acetylglucoside) increased over time, mirroring the development of red spots. While flavonoid identities were confirmed via the GNPS library, the anthocyanin structure was predicted using the *in-silico* tool SIRIUS. Due to the lack of a commercial standard for this specific predicted compound, experimental confirmation was not possible. Nevertheless, the combined transcriptomic and metabolomic patterns indicate a metabolic reprogramming that directs flux toward flavonoid and anthocyanin biosynthesis at infection sites, strongly linking the red pigment to anthocyanin accumulation.

Since red colouration can result from either anthocyanins or condensed tannins (proanthocyanidins, PAs), we used DMACA (p-dimethylaminocinnamaldehyde) stain to differentiate them. This stain specifically reacts with PAs to produce a blue-purple colour and was validated using a freshly cut apple as a positive control. The red pigments in the infected petals did not react with DMACA (Supplementary Figure S1), arguing against the presence of PAs. This finding, combined with other data, provides compelling evidence for anthocyanins. The LC-MS/MS identification of cyanidin 3-glucoside-5-(6-acetylglucoside), the petal-specific induction of anthocyanin genes, and the characteristic red-channel autofluorescence (600–650 nm) all strongly support that cyanidin-based anthocyanins is the primary source of the red colour. To truly confirm this compound’s structure, additional structural proof is required, e.g., by doing NMR on the purified compound. However, since we study a single cell response and the flower petals are just a few millimetres in size, this is technically not feasible. In the future, an alternative validation approach would be to chemically synthesise cyanidin 3-glucoside-5-(6-acetylglucoside) and match its LC-MS/MS characteristics to the experimental data.

Co-expression of flavonoid/anthocyanin biosynthetic genes with several MYB, bHLH and WD40 candidates in infected petals is consistent with the involvement of an MBW-type module in regulation of these compounds. Notably, the expression of the *C. seticuspe* genes that are orthologs to *Arabidopsis thaliana* bHLH TFs TT8/MYC1 are higher in mock-treated than infected *C. seticuspe* petals, making it unlikely that these operate as regulators of the induced program. This pattern points to either (i) recruitment of alternative MYB/bHLH partners during defence, (ii) a shift from developmental to stress-responsive regulators, or (iii) regulation that is not transcript-limited (e.g., pre-existing proteins, post-translational control, or cell-type effects). Discriminating among these possibilities will require targeted perturbation of the induced MYB/bHLH candidates and finer time-course profiling.

Re-activation of the anthocyanin pathway during *B. cinerea* infection can sometimes circumvent genetic blocks through activation of distinct regulatory nodes. In white *Antirrhinum majus* corolla tubes (the delila or del mutant), the anthocyanin pathway is normally blocked by absence of the bHLH transcription factor which regulates the expression of DFR. However, infection with *B. cinerea* stimulates the biosynthetic machinery downstream of the block, thereby restoring pigmentation (Harrison & Stickland, 1980). A similar phenomenon is observed in white-skinned grapes, where the canonical VvMYBA1 and VvMYBA2 transcription factors are silenced in those grapes. Notably, Blanco-Ulate et al. (2015) found that while these master regulators and the canonical VvUF3GT gene are not expressed during infection, a suite of alternative putative UF3GT genes were significantly up-regulated. This suggests that interactions with a fungal pathogen can activate independent regulatory nodes or alternative candidate genes to bypass primary genetic blocks, a mechanism that may similarly be at play in *C. seticuspe*.

Our data show a broad and coordinated induction of the flavonoid pathway in infected petals. Key biosynthetic genes, including CHS, CHI, F3H, DFR, arGST and UFGT were upregulated in concert with a significant accumulation of downstream metabolites such as quercetin, hyperoside and cyanidin-glucoside. This strong gene-to-metabolite correlation supports a major redirection of metabolic flux toward flavonoid and anthocyanin production at infection sites. A notable exception was the expression of the gene anthocyanidin synthase (ANS), for which we detected no transcripts. This could be due to several factors: i) expression may be highly transient and peak outside our sampling window; ii) there might be other ANS gene copies that are not in the genome annotation; or iii) regulation may occur post-transcriptionally, allowing for sufficient enzyme activity. While resolving the mechanism of ANS regulation requires further investigation, the consistent induction of genes upstream and downstream of this step, combined with the accumulation of the final cyanidin 3-glucoside-5-(6-acetylglucoside) provides compelling evidence that petals channel metabolic flux through the anthocyanin pathway during infection.

To validate the defensive function of the accumulated metabolites, we demonstrated their antifungal activity against *B. cinerea*. *In-vitro* assays revealed that infection-induced flavonoids (quercetin, hyperoside, acacetin) and anthocyanins (cyanidin derivatives) exhibited potent, dose-dependent inhibition of fungal growth. The observation that the cyanidin aglycone displayed higher antifungal potency than its glycosylated counterpart, cyanidin 3,5-diglucoside, suggests a bio-activation mechanism during infection. While the cyanidin glycoside appears less toxic *in-vitro*, it likely functions as a’concealed arsenal’ *in-vivo*. Fungal pathogens, including *B. cinerea*, secrete β-glucosidases as part of their pathogenicity strategy to degrade carbohydrates during infection. The *B. cinerea* genome contains around 40 genes that encode CAZymes with potential β-glucosidase activity (Drula et al., 2022). Such enzyme activity may strip the sugar moiety from stored, stable flavonoid glycosides, which are less active in fungal growth inhibition, and thereby release the less stable but more toxic aglycones at the site of infection. This mechanism has been demonstrated in mango-anthracnose interactions, where the release of aglycones such as cyanidin and quercetin by fungal β-glucosidase is more toxic than its glycosylated forms (Sudheeran et al., 2020). Thus, the accumulation of cyanidin glycosides in chrysanthemum petals may a stable pro-drug that is converted into more toxic compound after the release of fungal β-glucosidases. It is currently unclear which of the *B. cinerea* CAZymes would be involved in such a process.

Beyond direct toxicity of flavonoids and anthocyanins, they are also known antioxidants capable of scavenging reactive oxygen species (ROS), thereby mitigating oxidative stress at the infection site and preventing uncontrolled necrosis (Heller & Tudzynski, 2011). This dual capacity as both antimicrobial and antioxidant agents positions them as multifunctional protectants in the floral defence. Our observations are consistent with earlier reports that genetically modified anthocyanin-rich purple tomatoes are more resistant to *B. cinerea* than red tomatoes with no anthocyanins (Zhang, et al., 2013). Although our assays show clear antifungal activity for several infection-induced flavonoids and anthocyanins, it is likely that additional classes of secondary metabolites also contribute synergistically towards resistance against *B. cinerea*. In particular, terpenoids and other specialised metabolites may play equally important roles that were not studied here. Future work should therefore examine these additional pathways, especially the numerous biosynthetic gene clusters predicted by PlantiSMASH that were up-regulated after infection. Characterising the metabolites produced by these clusters may reveal previously unrecognised chemical defences deployed in infected flowers.

We leveraged the paired multi-omics dataset to identify candidate biosynthetic pathways using MEANtools, a dedicated computational framework for predicting plant specialised-metabolism pathways from integrated transcriptomics and metabolomics data. As one of the few tools specifically developed for multi-omics-driven pathway discovery in plants, MEANtools enable systematic prioritisation of transcript-metabolite associations and reconstruction of putative biosynthetic routes. Applying MEANtools to our dataset allowed to explore pathway-level organisation and identify potential enzymatic steps underlying the observed metabolic changes. This analysis identified a large anthocyanin-associated module that is presumably involved in the synthesis of potential phytoalexins, including mycosinol, as well as flavones such as chrysoeriol and selgin. However, this approach comes with some limitations. As the dataset is based on untargeted LC-MS/MS, MS² spectra were not available for several key metabolites identified by MEANtools such as the putative mycosinol feature (ID1317) and selgin (ID826) preventing confident *in-silico* structural annotation. MEANtools relies primarily on accurate mass and LOTUS database matches, which can lead to conflicts when MS² data are considered. For example, ID2951 was annotated by MEANtools as a compound similar to mycosinol but SIRIUS predicted a different sesquiterpenoid structure, and ID3945 predicted by MEANtools as chrysoeriol matched to phytosphingosine (a fatty-acid-derived lipid) in the GNPS library. Despite these constraints, the transcript–metabolite co-regulation within the module provides strong support for the occurrence of mycosinol-related defence response. Future targeted LC-MS/MS analysis with full fragmentation data would be essential to validate these predicted metabolites and determine whether mycosinol contributes to resistance of chrysanthemum against *B. cinerea*.

Taken together, our findings show that *Chrysanthemum seticuspe* flowers mount a multilayered chemical defence against *B. cinerea*, characterised by the transcriptional activation of the anthocyanin pathway and the accumulation of a broad suite of specialised metabolites, including candidate polyyne-type phytoalexins. The infection-triggered pigmentation response is not unique to *Chrysanthemum*, but reflects a broader pattern observed across angiosperms, in which normally white tissues such as white-skinned grapes, white-stage strawberry fruit, or the white corolla tubes of *Antirrhinum majus* produce localised red pigmentation (primarily anthocyanins) when infected by *B. cinerea* (Blanco-Ulate et al., 2015; Xu et al., 2025; Harrison & Stickland, 1980). Across these systems, stress-induced flavonoid and anthocyanin accumulation correlates with increased resistance. It is unclear whether this infection-triggered metabolic response is specific to necrotrophic pathogens or is also induced by pathogens with biotrophic or hemi-biotrophic lifestyle. It is also unknown whether comparable responses occur in cultivated *Chrysanthemum morifolium*, and whether their magnitude and composition differ between pigmented and non-pigmented cultivars. Future efforts combining targeted metabolite characterisation, functional genetics and high-resolution spatial/single cell-omics will be essential to uncover pathogen-induced plant defence metabolites within these floral tissues. Collectively, these studies open new avenues of research into floral defense mechanisms and the role of anthocyanin compounds in the interaction with necrotrophic pathogens, highlighting their functionality beyond their traditional ornamental roles.

## Acknowledgements

We thank all members of the Phytopathology Laboratory of WUR for technical assistance and helpful discussions. We acknowledge the support from Dr. Ric de Vos (Wageningen University & Research, The Netherlands) in the metabolomics work. We are grateful to Dr. Boas Pucker (University of Bonn, Germany), Dr. Nick Albert (Plant & Food Research, New Zealand) and Dr. Cathie Martin (John Innes Centre, UK) for their valuable scientific input and to Felicia Wolters (Wageningen University & Research, The Netherlands) for metabolomics data assistance and support. HS acknowledges funding from the TKI Graduate School Green Top Sectors, The Netherlands (grant LWV21.291, TU-2021-20). We thank Aike Post (Deliflor Chrysanten) and Dr. Nick de Vetten (Dekker Chrysanten) for fruitful discussions during project meetings, and we acknowledge Deliflor Chrysanten and Dekker Chrysanten for the supply of plant materials and for co-funding the project together with the TKI grant.

## Author contributions

HS contributed to conceptualisation, investigation, data analysis, visualisation, and writing—original draft, review, and editing. LR contributed to investigation, data analysis, visualisation, and writing—review and editing. FP contributed to data analysis and writing—review and editing. MS contributed to conceptualisation, methodology, visualisation, and writing—review and editing. KS contributed to methodology, data analysis, manuscript drafting, and writing—review and editing. JJJvdH contributed to methodology, supervision, manuscript drafting, and writing—review and editing. AF contributed to supervision and writing—review and editing. JK contributed to conceptualisation, methodology, visualisation, project administration, supervision, and writing—review and editing.

## Supplementary figures

**Figure S1:**
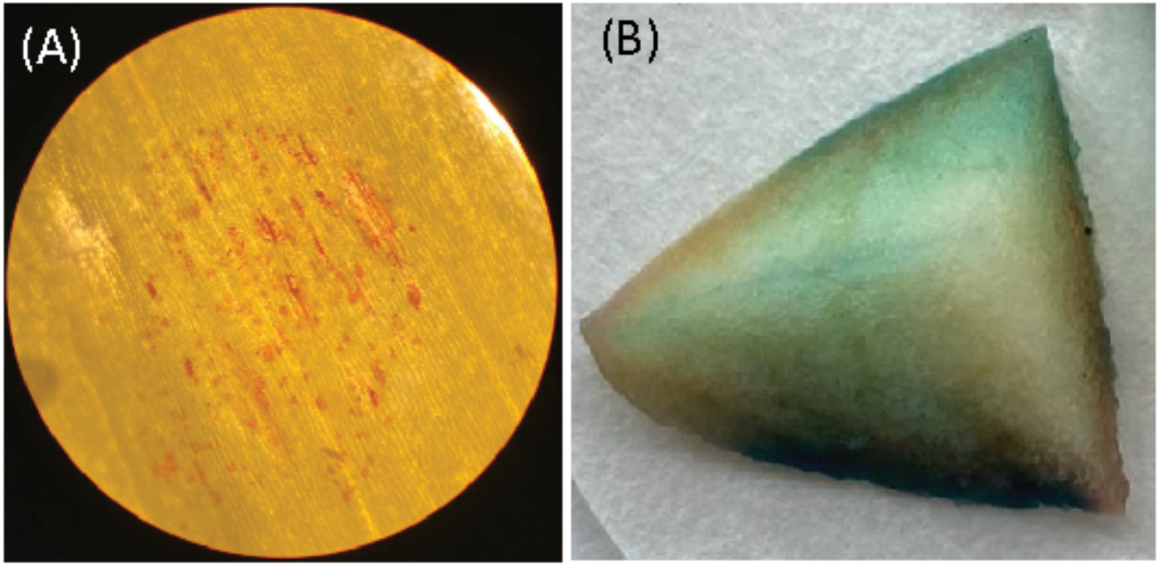
Figure A and B are DMACA staining of infected flower petals and freshly cut apples.

**Figure S2:**
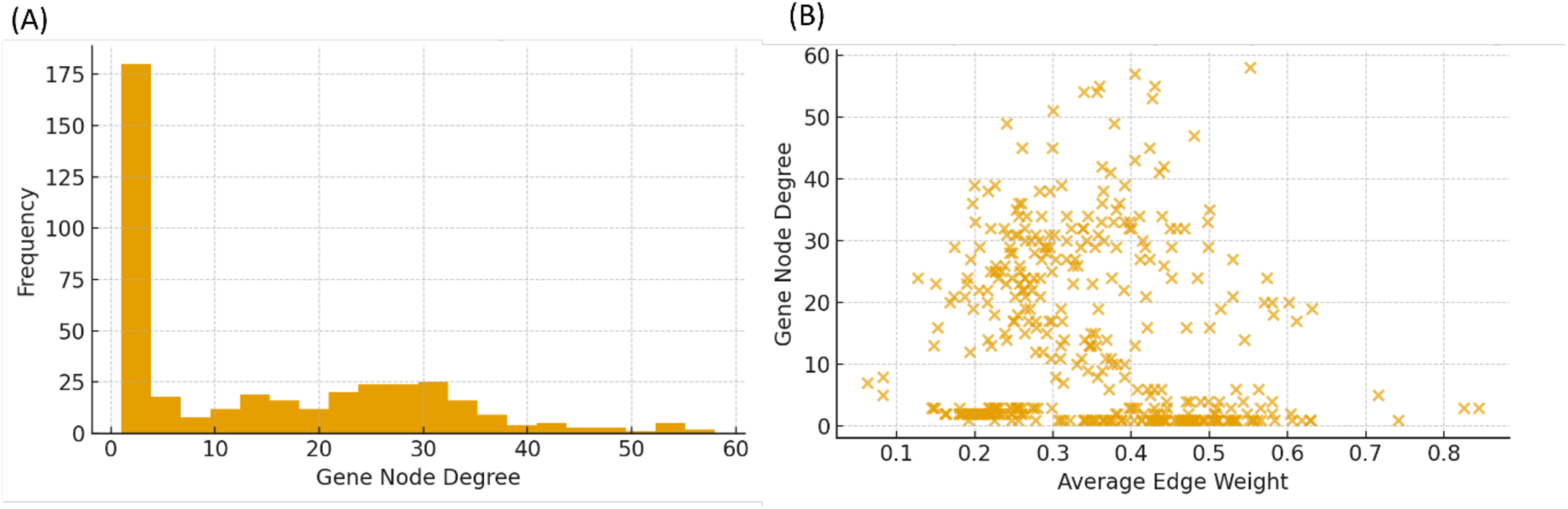
A) MEANtools predicted relationship between gene node degree vs frequency. B) MEANtools predicted relationship between gene node degree vs average edge weight.

**Figure S3:**
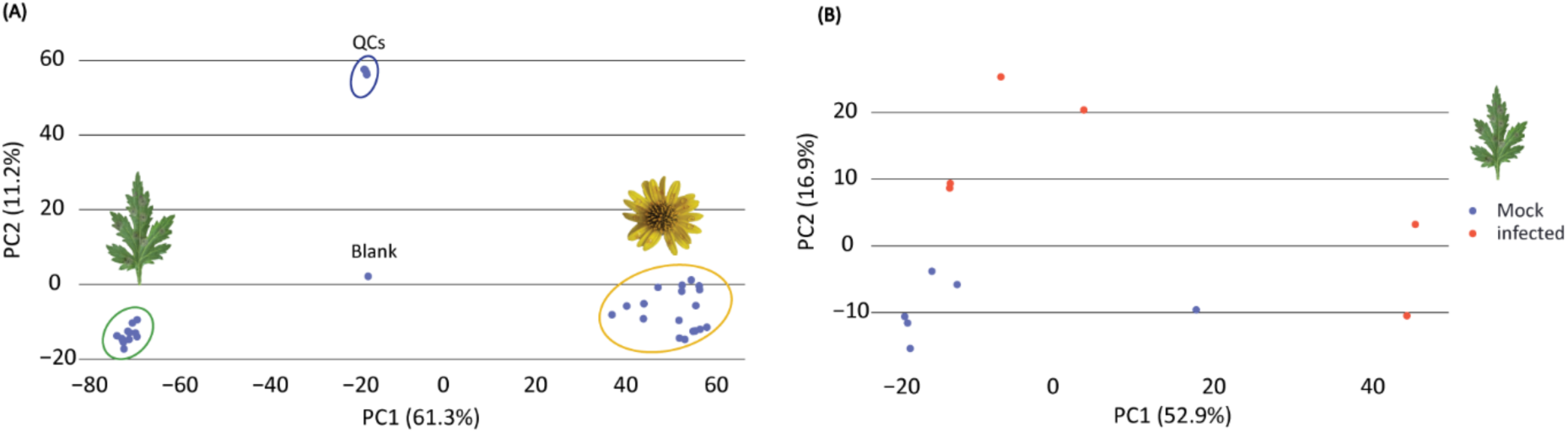
A) PCA of metabolite features from mock-inoculated, *B. cinerea*-infected samples, QCs and blank. B) PCA of metabolite features from mock-inoculated (blue) and *B. cinerea*-infected (red) leaf samples.

